# A Wearable Spiral Ultrasound Patch for Focused Ultrasound Peripheral Neuromodulation of Diabetic Neuropathic Pain

**DOI:** 10.1101/2024.12.12.628265

**Authors:** Cong Pu, Ben Fu, Pengda Lu, Xin Guan, Jiayi Zhang, Yuting Shen, Xiao Li, Lehang Guo, Huixiong Xu, Xiaoning Jiang, Chang Peng

## Abstract

Diabetic foot disease, a severe complication of diabetes, frequently results in diabetic neuropathic pain, which diminishes quality of life and increases the risk of amputation. Current treatment options are limited in their effectiveness and can present adverse effects. We have demonstrated a first-of-its-kind wearable spiral ultrasound patch for focused ultrasound peripheral neuromodulation to alleviate diabetic neuropathic pain. This innovative, stretchable ultrasound device reliably delivers focused ultrasound to peripheral nerves. The device’s ultrasound elements are arranged in a phyllotaxis spiral pattern, enhancing transmitted acoustic focus performance and minimizing grating and side lobes. Using a diabetic foot rat model, we confirmed peripheral neuropathy through behavioral assessments of mechanical allodynia and heat hyperalgesia, which showed significant pain relief after two weeks of ultrasound treatment. Additionally, nerve conduction velocity improved. Biological safety tests indicated no adverse effects on nerve tissues or cells. Our approach offers a non-invasive solution for managing diabetic neuropathic pain and provides critical insights into its underlying mechanisms, paving the way for more effective therapies.

## Introduction

Diabetic foot disease is one of the most serious complications of diabetes mellitus and a leading cause of amputation and mortality among diabetic patients [1, 2]. This condition results in significant suffering and financial costs for patients, while also imposing a heavy burden on their families, healthcare facilities, and society at large [3]. Key risk factors for developing diabetic foot ulcers, a chronic wound condition, include neuropathy, peripheral vascular disease, and decreased resistance to infections [4–6]. Among these, diabetic neuropathy (DN) is a primary contributor [4]. DN affects over 90% of diabetic patients, with at least 50% expected to develop DN over time [7]. This condition involves peripheral nerve dysfunction and loss of sensory function, typically starting in the lower extremities. The most common form, distal symmetric polyneuropathy, impacts the hands and lower limbs in a ‘stocking and glove’ pattern [7]. DN results in more hospitalizations than other diabetes-related complications and is the leading cause of non-traumatic amputations [8]. Moreover, research indicates a significant increase in the risk of diabetic foot ulceration due to DN [9]. Without adequate treatment, neuropathy can worsen, leading to a high risk of amputation (40% within 5 years) and a high mortality rate (up to 70% 5 years post-amputation) [7].

Diabetic neuropathy pain (DNP), a significant symptom of DN [10], impacting roughly 30%-50% of patients [11]. This pain is marked by sensations such as tingling, burning, sharpness, shooting pains, and electric shocks [3,8], often worsening at night and disrupting sleep. Conditions that affect the somatosensory nervous system can result in both loss of function and heightened sensitivity to painful stimuli (manifesting as hyperalgesia), allodynia, and spontaneous pain [11]. Consequently, DNP can severely diminish patients’ quality of life, daily activities, and mood, leading to social withdrawal and depression. Patients with DNP also have higher 10-year mortality rates compared to those without pain [11].

Managing DPN is challenging due to its complex pathophysiology and the limited effectiveness of existing pain relief treatments. Typically, treatment methods include glycemic control and the use of medications to alleviate pain [12]. However, most pharmacological treatments provide only symptomatic relief without addressing the underlying pathophysiological mechanisms. These treatments are also limited by side effects and the development of tolerance [10]. Although opioids can help control pain, they carry high risks of addiction, tolerance, and withdrawal symptoms upon discontinuation [13]. Both the American Academy of Neurology (AAN) and the Centers for Disease Control and Prevention (CDC) advise caution in using opioids for chronic, non-cancer pain, including DNP [7]. Opioids should be reserved for patients who do not respond to other treatments, as they are effective only for a small number of patients in the long term [11]. Interventional pain management techniques such as neural blockade, spinal cord stimulation (SCS), intrathecal medication and neurosurgical interventions are available [12], but these methods are often systemic and/or invasive. Some observational studies suggest that SCS may offer pain relief benefits, but the implantation process is generally considered too invasive [14].

Focused ultrasound is emerging as a promising non-invasive neuromodulation technique capable of precisely targeting and modulating neural activity. It offers a safe and accurate method of external regulation with various biological effects, including mechanical, cavitation, and thermal effects [15], making it superior to other strategies like light and magnetic field regulation, which often suffer from poor controllability, low precision and safety concerns [25]. Studies have demonstrated that specific ultrasound parameters can modulate a variety of ion channels on cell membranes, such as Piezo2 [16], the MS family [17], TRAAK+ [18], MscL [19], TRPA1 [20], and TRPV1 [21]. Ultrasound modulation can non-invasively activate specific neurons and trigger multiple physiological activities [16–18, 20], potentially blocking pain conduction.

Research has shown that high-intensity focused ultrasound (HIFU) can relieve pain, although the exact mechanism remains unclear. It is believed to involve localized denervation of tissue targets and/or neuromodulatory effects [22]. One notable feature of HIFU treatment is its rapid onset of pain relief, typically occurring within a day, and its long-lasting effects, with many patients experiencing benefits for up to a year. The increase in temperature induced by ultrasound is one mechanism that inhibits nerve conduction. Experiments have demonstrated that HIFU can safely and reversibly block nerve conduction in diabetic rats, offering potential peripheral pain relief [23]. Additionally, HIFU has been shown to increase both innocuous and noxious mechanical thresholds, as well as thermal withdrawal thresholds, in rodent models of diabetes [24].

Despite its significant potential for pain relief, current focused ultrasound transducers are bulky and rigid, making them unsuitable for daily wear and limiting their potential for long-term use [25]. Recent advancements in flexible electronics have paved the way for the development of wearable ultrasound devices for everyday healthcare applications [26–35]. In addition, various conformal ultrasound patches and stretchable ultrasound facial masks have recently emerged for therapeutic purposes [36, 37]. These devices primarily aim to stimulate the skin’s superficial surface but often neglect the need to target deeper nerves and optimize acoustic focus performance. For therapeutic applications targeting peripheral nerves located at specific depths beneath the skin, achieving precise focus performance is crucial. Thus, the development of a miniaturized ultrasound patch integrating portable wearability and therapeutic functions holds great promise for optimizing diabetic neuropathic pain relief. However, wearable ultrasound patches have yet to be reported for diabetic neuropathic pain relief.

In this study, we introduce a groundbreaking wearable spiral ultrasound patch specifically designed for focused ultrasound peripheral neuromodulation to alleviate diabetic neuropathic pain. This innovative, stretchable ultrasound device can reliably deliver focused ultrasound to the peripheral nerves. The ultrasound elements are arranged in a phyllotaxis spiral pattern, which enhances transmitted acoustic focus performance and minimizes grating and side lobes. A diabetic foot rat model was developed to validate the device, with peripheral neuropathy confirmed through behavioral tests. Pain was quantified using tests for mechanical allodynia and heat hyperalgesia. This approach not only provides deeper insights into the mechanisms underlying diabetic neuropathic pain but also paves the way for the development of new therapeutic interventions.

## Results

### Design of the wearable spiral ultrasound patch

Fig. 1a shows a wearable spiral ultrasound patch (w-SUP) attached to the shaved area of a rat with a diabetic foot ulcer. The patch is centered above the sciatic nerve to alleviate the rat’s diabetic neuropathic pain (DNP) by modulating the nerve through the thermal effect at the focal point of the ultrasound beam. The exploded view of the w-SUP, as illustrated in Fig. 1b, reveals its components: piezoelectric lead zirconate titanate ceramic (PZT) elements, vertical interconnect accesses (VIAs, Esolder-3022), flexible Cu/PI electrodes, and a flexible substrate (Ecoflex-0030). The w-SUP can conform to both spherical and cylindrical surfaces, as demonstrated in Fig. 1c and d, and can be stretched up to 60% strain from the result of the cyclic tensile test (Supplementary Fig. 1), showcasing its excellent mechanical compliance. E-solder 3022 paste was used to bond the stretchable electrodes to the transducer elements, with adhesion details shown in Fig. 1e,f and g. The overall dimensions of the w-SUP are 3.5 cm ×4 cm ×1.5 mm, and its total weight is just 9.05 g.

**Fig. 1.**
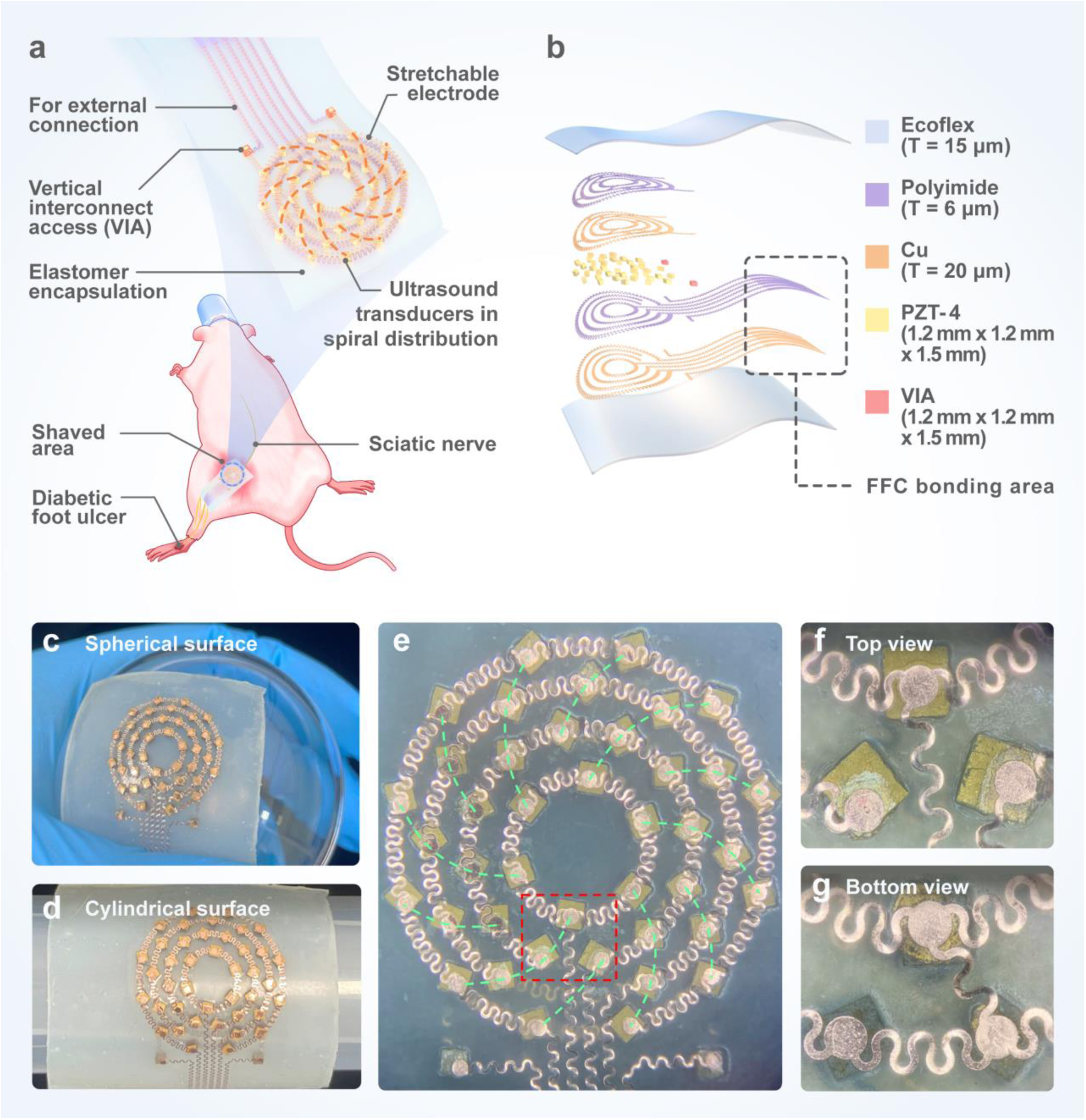
Overview of the wearable spiral ultrasound patch (w-SUP). **a** Schematic of the w-SUP conforming to the shaved area of a rat, with the center of the patch positioned above the sciatic nerve. **b** Schematic of the exploded view showing the components in the w-SUP, including Ecoflex, polyimide (PI), copper electrode, PZT-4 and vertical interconnect acess (VIA). **c**, **d** Optical images of the w-SUP conforming to spherical and cylindrical surfaces. Scale bar: 5 mm. **e** Optical image of the transducer elements with electrode adhesion. Scale bar: 1mm. **f**, **g** Enlarged images showing three elements within the red box from top and bottom views. Scale bar: 500 μm.

The designed transducer array comprises 41 square PZT-4 elements, each with dimensions of 1.2 mm × 1.2 mm × 1.5 mm. The transducer is designed to operate at a frequency of 1.5 MHz, with each element’s thickness nearly matching its half-wavelength (given the sound speed of PZT-4 is 4630 m/s, resulting in a half-wavelength of 1.54 mm).

### Acoustic performance of the w-SUP

The transducer elements are arranged in a phyllotaxis spiral pattern across four concentric circles within the patch, as illustrated in Fig. 2a, forming the four channels of the transducer rings. The overall distribution of the elements is confined to a 2 cm ×2 cm area, suitable for conforming to a rat’s skin. Utilizing a spiral structure disrupts the symmetry of the ultrasound array, thereby enhancing transmitted acoustic focus performance and reducing grating and side lobes effects [38–41]. In simulations with phase control, arranging the transducer elements in a phyllotaxis spiral pattern significantly reduces grating lobes compared to conventional 2D arrays (Supplementary Fig. 2). Additionally, even without phase control, the transducer exhibits natural focusing abilities in both circular and spiral patterns, unlike conventional 2D arrays (Supplementary Fig. 2).

**Fig. 2.**
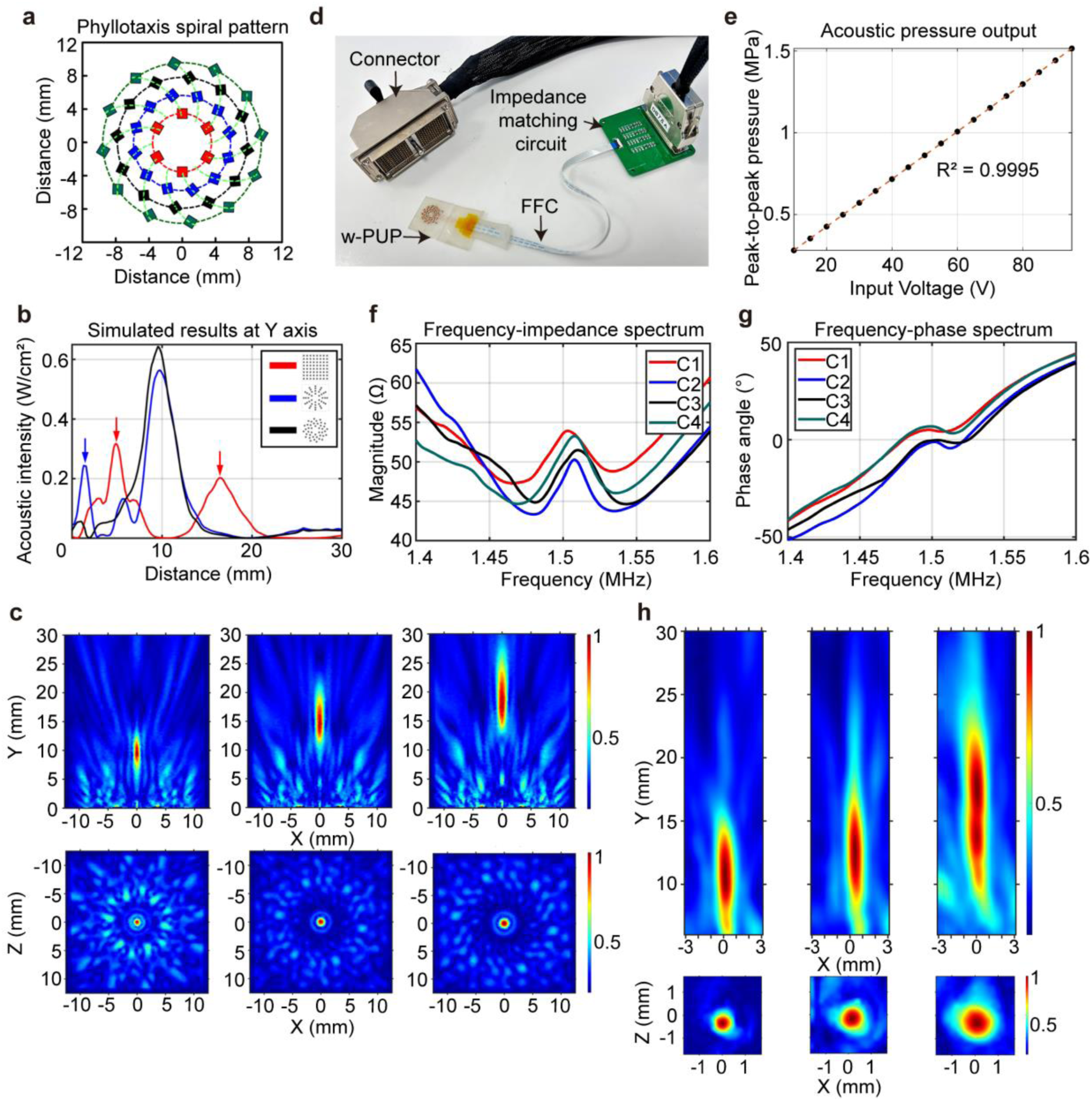
Simulation and characterization of the w-SUP. **a** Schematic diagram of the transducer elements arranged in a phyllotaxis spiral pattern. **b** Simulated acoustic intensities for three different patterns: red for the square pattern, blue for the circular pattern, and black for the phyllotaxis spiral pattern. **c** Simulated acoustic beam profiles in the Y-X axis and Z-X axis of the phyllotaxis spiral pattern without phase control, with targeted focal lengths of 14 mm and 20 mm, respectively, from left to right. **d** Optical image of the ultrasound patch and its electrical connections, including a flexible flat cable (FFC), an impedance mathing circuit and an ITT connector. **e** Measured acoustic pressure output at the focal point of the w-SUP as a function of input voltage (peak-to-peak). **f, g** Frequency-impedance and frequency-phase spectra of the four channels of the w-SUP after impedance matching. C1 to C4 correspond to the four channels of the w-SUP. **h** Measured acoustic beam profiles in the Y-X axis and Z-X axis of phyllotaxis spiral pattern without phase control, with targeted focal lengths of 14 mm and 20 mm, respectively, from left to right.

In the simulated acoustic intensity results shown in Fig. 2b, without phase control, the maximum acoustic intensities from the transducer in the spiral, circular and square patterns are 0.63 W/cm^2^, 0.58 W/cm^2^, and 0.34 W/cm^2^, respectively. The ratio of the main lobe intensity to the grating lobe intensity in the spiral, circular and square patterns are 10.36 dB, 3.06 dB, and 1.52 dB, respectively. As illustrated in Fig. 2c, the simulated focal lengths of transducer in the spiral pattern are 9.5 mm (without phase control), and 14.7 mm, and 17.6 mm when the targeted focal lengths are 14 mm and 20 mm, respectively. Furthermore, the acoustic pressure at the focal point of transducer in the spiral pattern surpasses that in the circular pattern, both with and without phase-control focusing (Supplementary Tab. 1). The beam profiles of the ultrasound transducers arranged in the spiral pattern were simulated at frequencies ranging from 0.5 MHz to 2 MHz (Supplementary Fig. 2 and 3). Notably, the transducer operating at 1.5 MHz has a natural focal length of 9.5 mm, which is approximately the distance between the sciatic nerve of a rat and its skin (10 mm). Additionally, the simulated acoustic pressure at the focal point is the highest with and without phase control (Supplementary Tab. 1).

The w-SUP was electrically connected to a Vantage HIFU system (Verasonics, USA) via a flexible flat cable (FFC), an impedance matching circuit, and a custom DL5-260P ITT connector, as illustrated in Fig. 2d. Before impedance matching, the average resonance frequency, antiresonance frequency, impedance, and phase angle at resonance frequency of the four channels were 1.49 MHz, 1.53 MHz, 2040 Ω, and - 9.28°, respectively (Supplementary Fig. 4). After impedance matching, these values were adjusted to 1.47 MHz, 1.51 MHz, 45.03 Ω, and −9.68°, respectively (Fig. 2f and g).

Fig. 2e illustrates the correlation between the measured acoustic pressure output at the focal point and the input voltage. The curve, derived from 18 data points, has a slope of 0.0145 and an intercept of 0.1354, with a coefficient of determination (R²) of 0.9995. Furthermore, the maximum peak-to-peak acoustic pressure of the phyllotaxis spiral array was evaluated on two different curved surfaces (Supplementary Fig. 5). The peak-to-peak acoustic pressures at the focal point of the w-SUP adhered to a flat plane, a 20°curved surface, and a 40°curved surface were 0.39 MPa, 0.45 MPa, and 0.44 MPa, respectively. The measured focal lengths of the w-SUP adhered to a flat plane were 11 mm (without phase control), 12.7 mm, and 17.5 mm for targeted focal lengths of 14 mm, and 20 mm, respectively, as shown in Fig. 2h. The measured full width at half maxima (FWHM) in the Y-X and Z-X planes are detailed in Supplementary Tab. 2.

### Thermal performance of the w-SUP

As depicted in Fig. 3a, the sciatic nerve is positioned roughly 10 mm beneath the skin and has an approximate thickness of 2 mm, as determined from the ultrasound image. During ultrasound therapy, the ultrasound beam is focused on the nerve by aligning the center of an ultrasound imaging probe with the center of the ultrasound patch. Fig. 3b displays the front and top views of the w-SUP adhered to a rat’s skin. The radius of curvature for the w-SUP on the rat’s skin is 18.34 cm (Supplementary Fig. 6). Fig. 3c illustrates the acoustic beam profiles without phase control for the w-SUP on a 20°curved surface (with a radius of curvature of 18.34 cm) and a 40°curved surface (with a radius of curvature of 9.17 cm), showing focal lengths of 10.5 mm and 8.7 mm, respectively. The temperature profiles, measured after 5 min of sonication for the w-SUP on both the 20°and 40°curved surfaces, are shown in Fig. 3d-i. Temperature changes were determined by subtracting 37°C from the readings recorded by a thermocouple in pork tissue.By varying input voltage, duty cycle and pulse cycle number, the ultrasound sequence with the highest heating effect within 5 min was identified. Specifically, with an input voltage of 600 mVpp, an 80% duty cycle, and 20 cycles, the w-SUP on the 20°and 40°curved surfaces achieved the highest temperature rises of 6.2°C and 4.5°C at the focal spot, respectively.

**Fig. 3.**
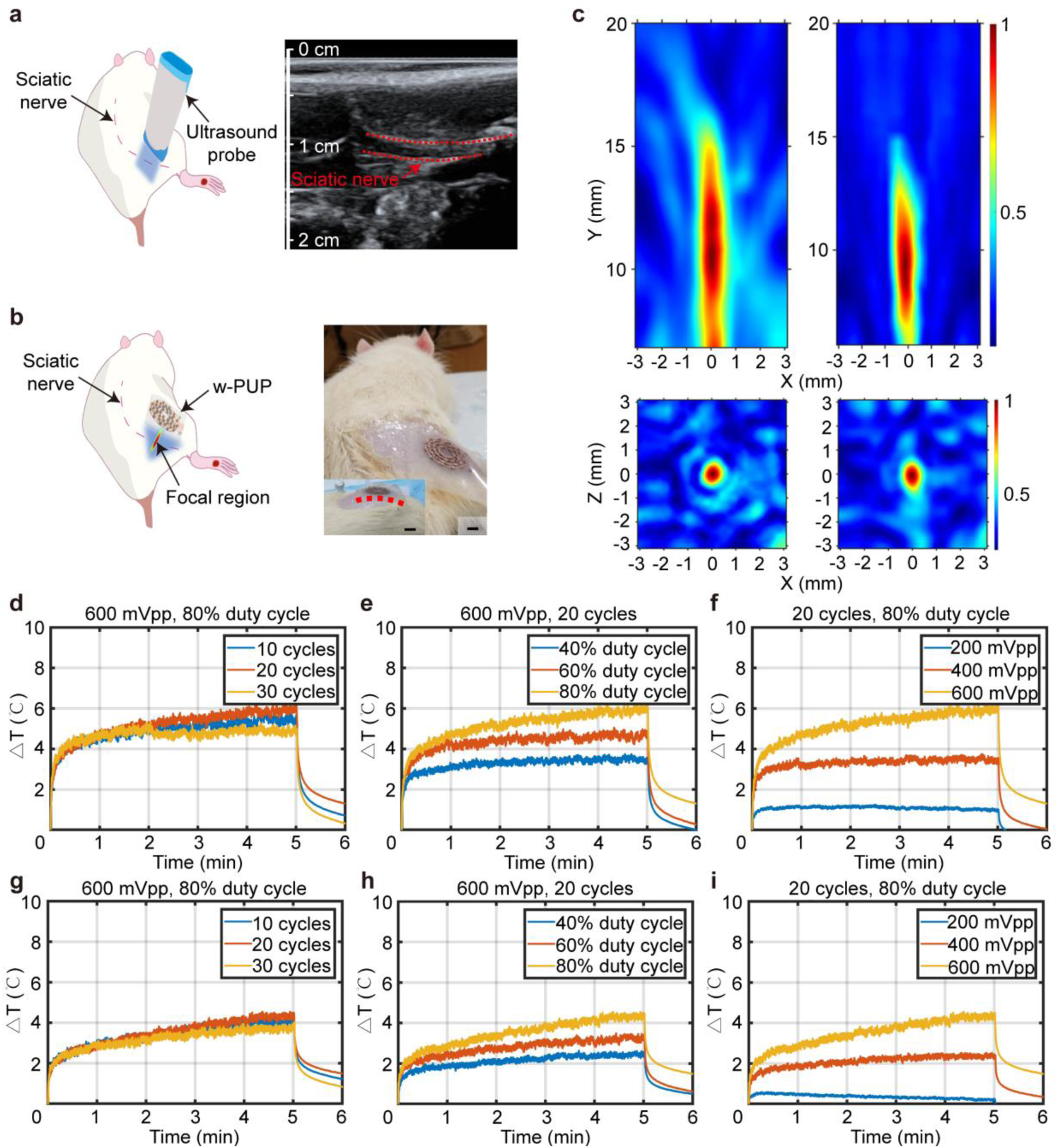
Demonstration of temperature rising capability using the w-SUP. **a** Schematic of sciatic nerve imaging in a rat, with the the sciatic nerve indicated by red dashed lines. **b** Schematic of the w-SUP modulating the sciatic nerve, along with optical images showing the front view of the w-SUP attached to the shaved area of the rat. The inset shows the top view. Scale bar: 5 mm in the front view and 1 cm in the top view. **c** Measured acoustic beam profiles of the w-SUP adhered to surfaces with 20°and a 40°curves, shown from left to right, respectively. **d, e, f** Measured temperature profiles for the w-SUP adhered to a 20°curved surface under three conditions: (left) with a fixed input voltage and duty cycle at three different cycle numbers, (middle) with a fixed input voltage and cycle number at three different duty cycles, and (right) with a fixed cycle number and duty cycle at three input voltages. **g, h, i** Measured temperature profiles for the w-SUP adhered to a 40°curved surface under the same three conditions as in **d**, **e**, and **f**.

### Variation in mechanical allodynia and heat hyperalgesia

The procedure for the animal experiments is illustrated in Fig. 4a. A total of 30 male Sprague Dawley (SD) rats were divided into five groups (further details provided in the Methods section). To assess the effects of ultrasound treatment on DNP, both behavioral and nerve conduction tests were conducted. The treatment protocols for the single treatment (ST) and multiple treatments (MT) groups are shown in Supplementary Fig. 7. In the ST group, DNP was quantified on Day 0 and Day 1, with the w-SUP treatment administered on Day 1. In the multiple treatments group (MT), DNP was quantified on Day 0, Day 7, and Day 14, with the w-SUP treatments administered daily from Day 1 to Day 14.

**Fig. 4.**
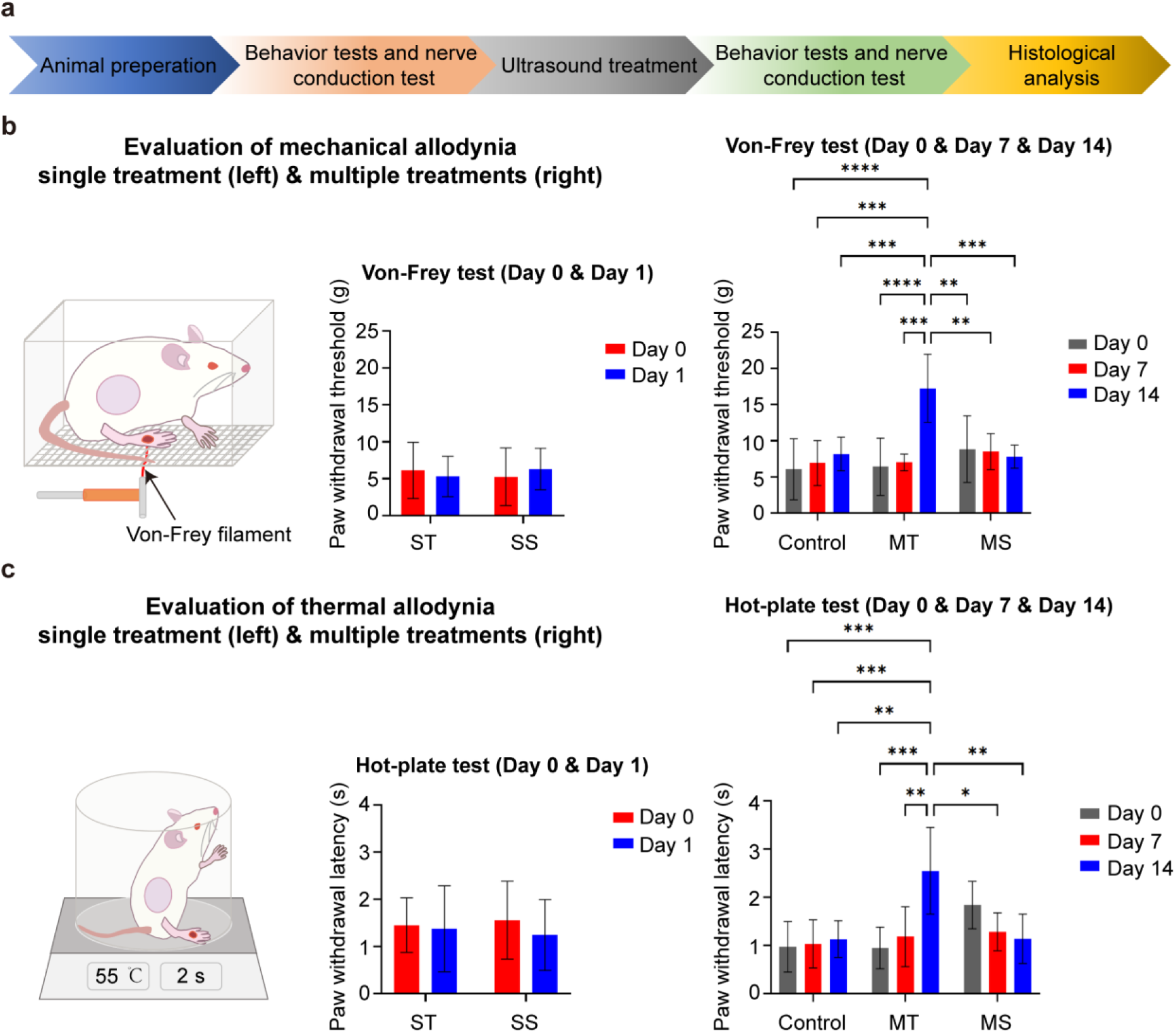
Animal experiments design and behavior tests of rats. **a** Procedure of animal experiments. **b** Schematic diagram for the evaluaiton of mechanical allodynia in rats. Results from the Von-Frey test for the single treatment (ST) group and the multiple treatments (MT) group. In the ST group, the F value is 0.4688 with a degree of freedom (DF) of 1. In the MT group, the F value is 6.357 with a DF of 4. **c** Schematic diagram for the evaluaiton of heat hyperalgesia in rats. Results from the Hot-plate test for the ST group and the MT group. In the ST group, the F value is 0.1416 with a DF of 1. In the MT group, the F value is 6.249 with a DF of 4. Stastical significance was determined by two-way ANOVA with Tukey’s multiple comparisons (n = 6, *p<0.05, **p<0.01, ***p<0.001, ****p<0.0001). Data are presented as mean ±standard deviations (SD), with error bars representing SD. SS: single sham, MS: multiple sham.

As illustrated in Fig. 4b, the paw withdrawal force for the ST group slightly decreased from Day 0 (6.13 ±3.80 g) to Day 1 (5.30 ±2.74 g), while the single sham (SS) group displayed an increase from 5.25±3.91 g to 6.30 ±2.81 g over the same period. No significant difference was found between the ST and SS groups regarding mechanical allodynia after one day of ultrasound treatment (p = 0.9545). After two weeks, the MT group demonstrated a significant increase in paw withdrawal force, rising from 6.40 ±3.95 g on Day 0 to 17.25 ±4.71 g on Day 14, indicating a strong response to treatment. In contrast, the control and multiple sham (MS) groups displayed more modest changes, with the MS group experiencing a slight decrease by Day 14. Significant differences were recorded between the MT and MS groups (p = 0.0005) and between the MT and control groups (p = 0.0009) by Day 14.

In Fig. 4c, paw withdrawal latency in the ST group remained relatively stable from Day 0 (1.45 ± 0.58 s) to Day 1 (1.38 ±0.91 s), while the SS group showed a slight decrease from 1.56 ±0.82 s to 1.25 ±0.75 s. No significant difference was observed in heat hyperalgesia between the ST and SS groups after one day of treatment (p = 0.9909). Over 14 days, the MT group exhibited a significant increase in paw withdrawal latency, from 0.95 ± 0.43 s on Day 0 to 2.55 ± 0.90 s on Day 14. Meanwhile, the control and MS groups showed minimal changes, with the MS group experiencing a slight decrease by Day 14. By Day 14, significant differences were noted between the MT and MS groups (p = 0.0013) and between the MT and control groups (p = 0.0012).

### Variation in nerve conduction velocity

As shown in Fig. 5a, a set of five electrodes is employed to record the compound action potential (CAP) in a rat model. The nerve conduction velocity is calculated by dividing the distance between the S1 and R2 electrodes by the latency of the CAP peak. An optical image of the nerve conduction test setup is provided in Fig. 5b. The representative CAP responses for a rat in the MT group on Day 0, Day 7 and Day 14 are shown in Fig. 5c.

**Fig. 5.**
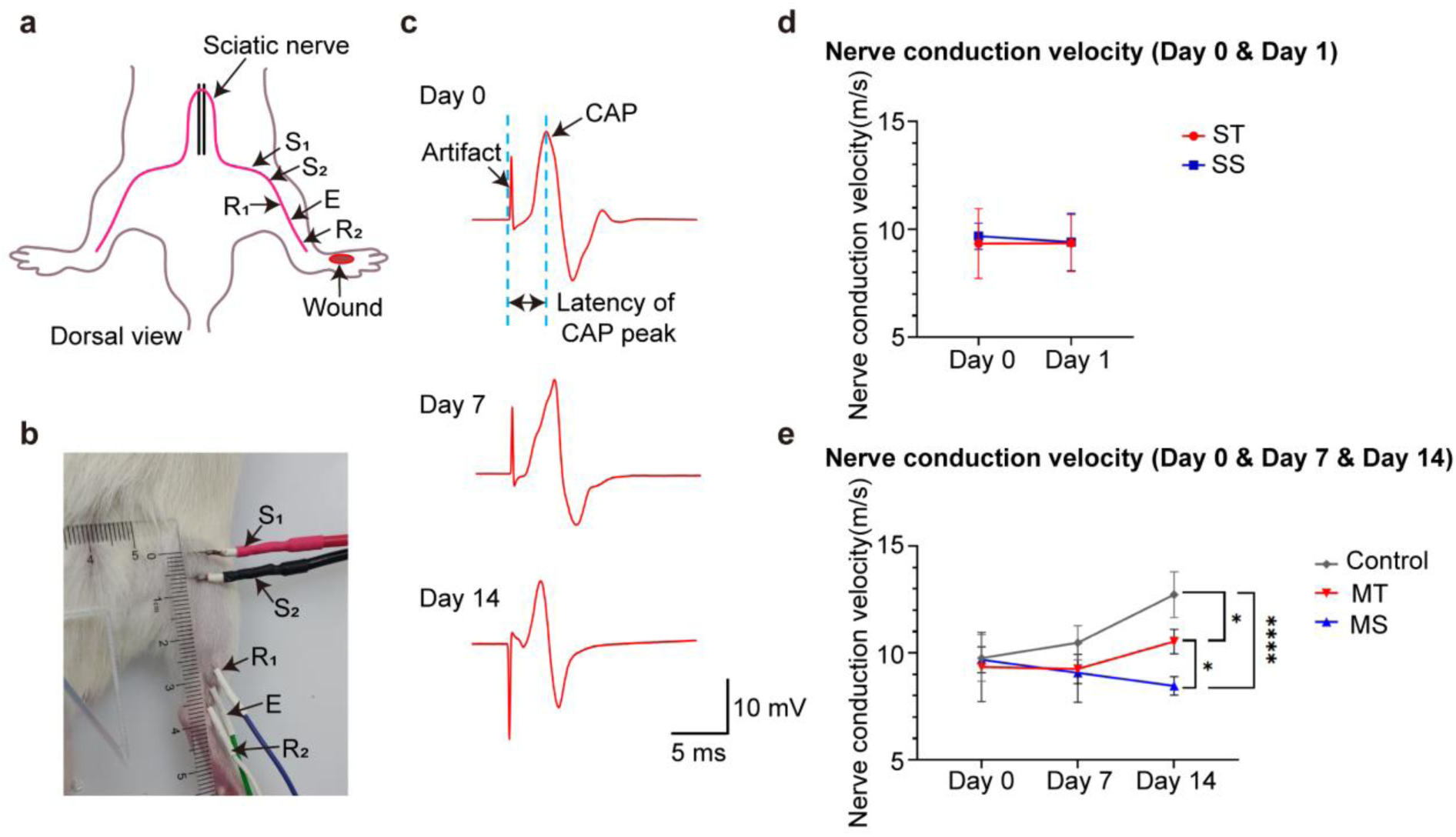
Nerve conduction test of rats. **a** Schematic diagram illustrating the positions of the electrodes for compound action potential (CAP) measurement, S1: stimulating electrode (+), S2: stimulating electrode (−), R1: recording electrode (+), R2: recording electrode (−) and E: reference electrode. **b** Optical image of the nerve conduction test being performed on a rat in the multiple treatments (MT) group. **c** Reprensentative CAP response results from a rat on Day 0, Day 7 and Day 14. **d** Results from the nerve conduction tests for both the single treatment group (ST) and the MT group. Stastical significance was determined by two-way ANOVA with Tukey’s multiple comparisons (n = 6, *p<0.05, **p<0.01, ***p<0.001, ****p<0.0001), Data are presented as mean ± standard deviations (SD), with error bars representing SD. SS: single sham, MS: multiple sham. For the ST group, F value is 0.1416 with a degree of freedom (DF) of 1. For the MT group, the F value is 7.112 with a DF of 4.

As depicted in Fig. 5d, the nerve conduction velocities for the ST group were recorded as 9.34 ± 1.62 m/s on Day 0 and 9.69 ±0.61 m/s on Day 1. For the single sham (SS) group, the velocies were 9.37 ±1.32 m/s on Day 0 and 9.41 ±0.75 m/s on Day 1. No significant differences were observed in nerve conduction changes between the ST and SS groups after one day of ultrasound treatment (p value is 0.9999).

After two weeks, as shown in Fig. 5e, the nerve conduction velocity in the control group exhibited a consistent increase from 9.76 ± 1.09 m/s on Day 0 to 12.73 ± 1.07 m/s on Day 14, indicating progressive nerve recovery. The MT group showed moderate improvement, with nerve conduction velocity increasing from 9.34 ±1.62 m/s on Day 0 to 10.53 ±0.57 m/s on Day 14. Conversely, the MS group displayed a slight decline, with the velocity decreasing from 9.68 ±0.61 m/s on Day 0 to 8.47 ±0.43 m/s on Day 14. On Day 14, there were significant differences between the MT and MS groups (p = 0.0013) as well as between the MT and control groups (p = 0.0012).

### Biological safety assessment of the w-SUP

Given the long-term nature of therapeutic ultrasound procedures, it is crucial to assess the biological safety of the w-SUP. As illustrated in Fig. 6a, cell staining results indicated that the survival fraction of PC12 cells exposed to focused ultrasound for 30 min remained unchanged. Fig. 6b illustrates the healing progression of diabetic foot ulcers in the control, MS and MT groups. After 14 days of ultrasound treatment, the MT group exhibited the most rapid wound recovery compared to the other groups. Additionally, Fig. 6c presents the H&E staining images, revealing that the sonication with the w-SUP appears to be safe, with no apparent damage observed when compared to the sham and control groups.

**Fig. 6.**
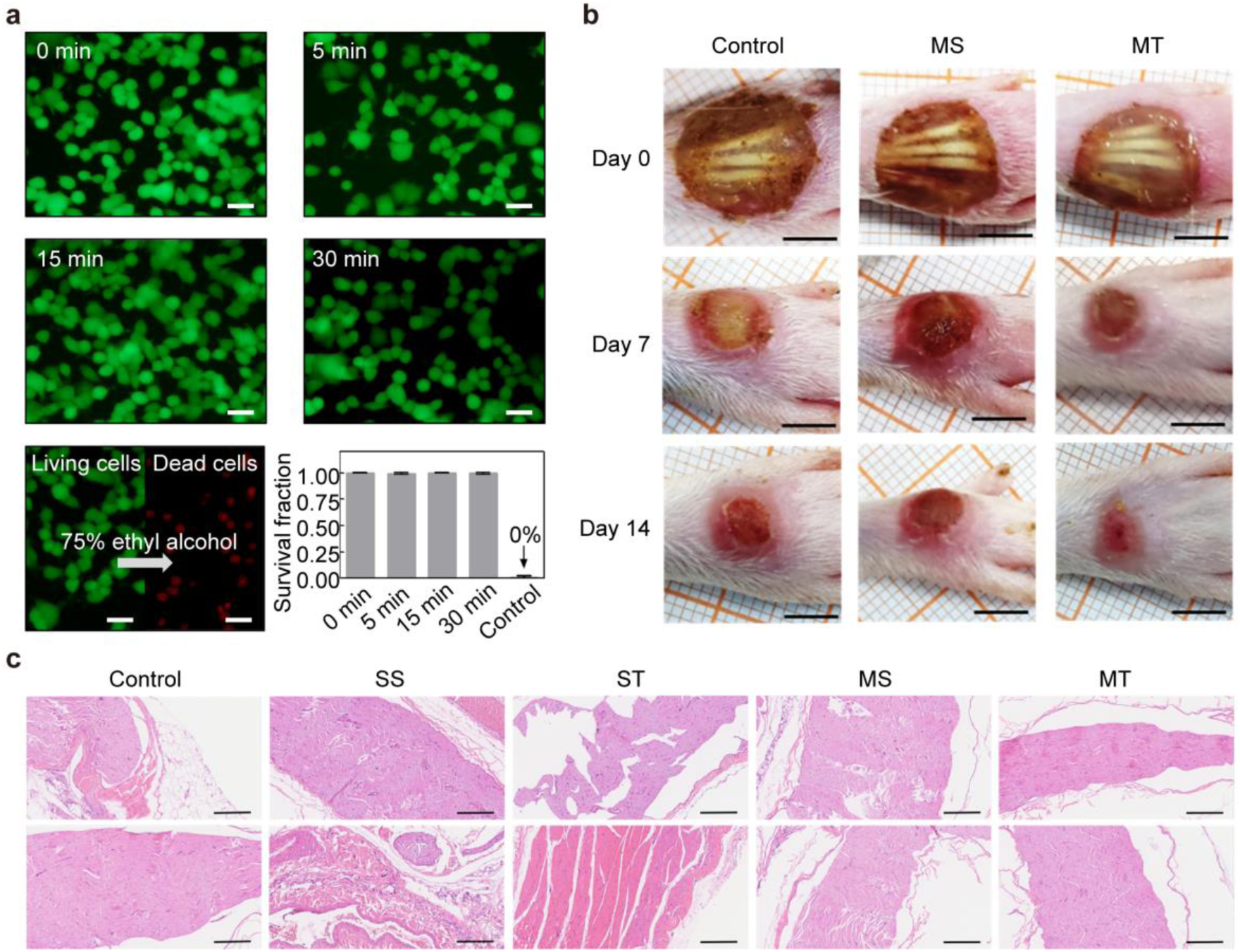
Biological safety evaluation of the w-SUP and observations of wound healing in rats. **a** Fluorescence images of live (green) and dead cells exposed to ultrasound for 0 min, 5 min, 15 min and 30 min. The control group consisted of cells treated with 75% ethanol for 10 min. Scale bars are 25 μm. **b** Traces of wound closure over 14 days among control, multiple sham (MS) and multiple treatment (MT) groups of rats. Scale bars are 5 mm. **c** H&E staining images of sciatic nerve sections cut in the sagittal plane. SS: single sham, ST: single treatment. Scale bars are 200 μm.

## Discussion

In this study, we developed a wearable phased array ultrasound patch (w-SUP) incorporating PZT-4 transducer elements, stretchable Cu/PI electrodes, and a flexible Ecoflex substrate. Designed for focused ultrasound (FUS) peripheral neuromodulation, the w-SUP aims to alleviate diabetic neuropathic pain (DNP). This patch consistently delivers FUS to peripheral nerves, effectively raising localized temperature to 43°C. Diabetic foot rat models with peripheral neuropathy and diabetic foot ulcers were established to evaluate its efficacy. Pain levels were assessed using behavioral tests including mechanical allodynia and heat hyperalgesia. After 14 days of FUS treatment, significant differences were observed between multiple treatment groups and control groups in both Von-Frey test and Hot-plate test. Additionally, nerve conduction velocity tests showed significant improvements in the MT group compared to both control group and MS group. The biological safety of the w-SUP was also confirmed through comprehensive tests.

In the acoustic simulation, transducer elements arranged in a phyllotaxis spiral pattern exhibited superior acoustic performance, demonstrating fewer grating lobes, natural focusing ability and greater acoustic intensity under the same driving voltages at 10 mm compared to other patterns. However, to simplify the manufacturing process, the 41 transducers are divided into 4 channels for connection instead of having 41 individual channels. This led to some differences between the simulated and measured beam profiles. By employing multi-layer electrode technology [29], each transducer element can be connected individually, enabling precise control to enhance focusing performance and adapt to irregular surface curvatures.

Our previous study demonstrated that pulsed stimulation provides superior temperature control and stability compared to continuous stimulation [42]. The thermal performance of the w-SUP was evaluated using pork tissue under different pulsed ultrasound sequences. Acoustic power and duty cycle were identified as the primary factors influencing the rate of temperature increase, while varying the cycle number from 20 to 30 had minimal impact on temperature variations.

The results of behavior tests in Fig. 4b and c indicated that mechanical allodynia and heat hyperalgesia in rats were not significantly improved after one or seven days of ultrasound treatment, likely due to the short recovery period following wound modeling. However, after two weeks of ultrasound treatment, significant reductions in pain sensation were observed, with mean values of PWT and PWL exceeding those of the other groups. It is suggested that focused ultrasound (FUS) therapy may enhance blood flow, thereby improving vascular supply to the endoneurium and nerve fibers, which could facilitate the repair of damaged nerve fibers and promote wound healing [43]. Furthermore, the decline in nerve conduction velocity observed in diabetic rats was mitigated following a two-week treatment with the w-SUP. This treatment might help reduce the degradation of the myelin sheath surface area, the myelin-to-axon ratio, and the number of unmyelinated axons or small fibers [44]. The potential improvement in microcirculation brought about by the ultrasound therapy could counteract the decrease in conduction velocity associated with DNP.

Safety is another crucial factor in evaluating the application potential of the w-SUP. The histological analyses and cell safety tests indicated that ultrasound stimulation from the w-SUP does not damage neural tissues or cells. Additionally, the *I_spta_, I_sppa_*, mechanical index (MI), and CEM43°C values for 1.5 MHz pulsed stimulation (600 mVpp, 80% duty cycle, 20 cycles and 5 min of sonication) are 208 mW/cm^2^, 639 mW/cm^2^, 0.59 and 1.35 min, respectively. Those values fall within the recommended safety limits (*I_spta_* < 720 mW/cm^2^, *I_sppa_* < 190 W/cm^2^, MI < 1.9 and CEM43°C < 2 min) [45, 46]. To assess the potential thermal impact of the w-SUP on the human body, skin temperature was monitored using a handheld thermal infrared (IR) imager during a 5-minute ultrasound application to the forearm. The recorded temperature variations before and after ultrasound exposure ranged from 0.3 to 0.4°C. No signs of discomfort, such as skin irritation or itching, were reported after 5, 10, or even 15 minutes of w-SUP use (Supplementary Fig. 8).

The results of the behavioral and nerve conduction tests are promising; however, a deeper understanding of how FUS interacts with cellular and physiological mechanisms of pain alleviation is essential. Clinical trials are necessary to evaluate the treatment’s effectiveness in patients with diabetic foot ulcers. While our study demonstrates that the w-SUP is both effective and safe, challenges remain in making it suitable for daily use. Further miniaturization of the electrical connection between the w-SUP and the ultrasound activation systems is required. We employed a 3D imaging system (as detailed in the Methods section) to accurately position the transducers, but this method is impractical for everyday applications. To address this, optical shape sensing fiber has been utilized to detect the real-time positions of each transducer element [47], which may facilitate the functionalization and miniaturization of the device.

## Methods

### Fabrication process of stretchable electrode

The fabrication process began with designing the stretchable electrode pattern by using AutoCAD software (Autodesk, USA). A polymide (PI) solution was spin-coated onto a Cu foil at 500 rpm for 30 s, then at 4000 rpm for 60 s. Simultaneously, polydimethylsiloxane (PDMS, mixed at a 20:1 ratio) was spin-coated onto a glass slide substrate at 1000 rpm for 30 s and then 2500 rpm for 60 s. The PDMS layer was cured in an oven at 45 °C for 6 h. The PI layer underwent a staged curing process on a hot plate: 80 °C for 10 min, 120 °C for 10 min, 180 °C for 20 min, 200 °C for 10 min, 220 °C for 10 min, and finally 250 °C for 10 min. The cured Cu/PI layer was then bonded to the PDMS-coated glass slide, completing the substrate preparation.

A laser ablation system (LPKF ProtoLaser U4, 2 W power, 80 kHz pulse repetition frequency) was employed to etch the electrode pattern onto the Cu/PI layer. Water-soluble tape was used to lift the top and bottom Cu/PI electrodes from their PDMS donor substrate. The receiver substrate was made of Eco-flex and a glass slide coated with Polymethylmethacrylate (PMMA). PMMA was spin-coated onto the glass slide at 500 rpm for 20 s and then at 1500 rpm for 60 s, followed by curing at 60 °C for 20 min. Subsequently, Eco-flex was spin-coated onto the PMMA at 1000 rpm for 60 s and cured at room temperature for 40 min. To enhance the bonding between the electrodes and Eco-flex, UV ozone surface activation was applied for 3 minutes. The water-soluble tape was dissolved by immersing it in hot water. The top and bottom electrodes are depicted in Supplementary Fig. 9, and simplified schematic diagrams of the fabrication process are shown in Supplementary Fig. 10.

### Assembly of piezoelectric elements and soft substrate

PZT-4 ceramic (Hongsheng Acoustics, China) was selected as the active material for its high-quality factor (Q) for high power transmit applications. The ceramic was polished to a thickness of 1.5 mm using a Precision Lapping and Polishing System (MCF, China). Subsequently, a Cr/Au (50/100 nm) electrode was sputtered onto the lapped surface by physical vapor deposition (Pro Line PVD 75, Kurt J. Lesker, USA). The PZT-4 material was then diced into the size of 1.2 mm × 1.2 mm using a wafer dicing saw (DAD323, DISCO, Japan). E-solder 3022 (Von Roll USA Inc., Schenectady, NY, USA) was used for vertical interconnect access (VIA). Two E-solder elements were fabricated via the same lapping and dicing process. A mold was used to maintain relative positions while 41 PZT-4 elements and 2 VIAs were welded onto the top electrode with E-solder paste, then cured in an oven at 45°C for 6 h. Anisotropic conductive film (ACF) tape was used to bond a flexible flat cable (FFC) to the top electrode. The bottom electrode was subsequently welded onto the transducers and VIAs using the same procedure. The entire assembly was encapsulated in Eco-flex, which was cured at room temperature for 2 hours. Finally, the glass slides were removed by dissolving the PMMA layer between the Eco-flex and the glass slide with acetone for 1 h.

### Electrical performance analysis and connectivity assessment

An impedance analyzer (E4990A, Keysight Technologies, USA) was utilized to measure the electrical impedance of the transducer elements arranged in four phyllotactic spiral lines. The ultrasound patch was electrically connected to a PCB with a series of capacitors and inductors, designed to eliminate the imaginary component of its impedance and match the real component to the desired 50 Ω. Subsequently, the PCB was connected to a Vantage HIFU system (Verasonics, USA) via a custom DL-260P ITT connector.

### Cyclic tensile tests

The tensile properties of the w-SUP was evaluated using cyclic tensile tests. These tests were conducted on a universal testing system (INSTRON) at a loading rate of 1 mm/s, with strain ranges varying from 10% to 60% and 10 cycles conducted per test.

### Finite element analysis

Commercial software COMSOL Multiphysics was employed for optimizing the array layout (Supplementary Fig. 2 and 3). The simulation primarily focused on acoustic beam profiles of transducer with different patterns and frequencies of the transducer elements. The physical fields considered in the simulation included pressure acoustics, solid mechanics and electrostatics. Additionally, the acoustic field was analyzed under both natural focusing conditions and phase control conditions.

### Animal preparations

All animal experiments were conducted in accordance with the Animal Care and Use Committee of ShanghaiTech University (approve number: 20231129001). Male Sprague Dawley (SD) rats (n=30) were randomly assigned into five groups, including control group (n=6), single treatment group (ST, n=6), single sham group (SS, n=6), multiple treatment group (MT, n=6), and multiple sham group (MS, n=6). All rats, except those in the control group, were intraperitoneally injected with 1% Streptozotocin (STZ) solution (dose: 45 mg/kg) after fasting one night in advance. Following the injection, the rats fasted for an additional 2 h before resuming their regular diet. Each rat was then anesthetized with 2% pentobarbital sodium (0.2 mL/100g), and a 1 cm diameter wound was created on the dorsum of the right hind paw using scissors.

### Assessment of ultrasound patch curvature on rat’s skin

The curvature of the ultrasound patch on a rat’s skin was characterized using an instantaneous 3D imaging system (VR3D Performance4D, Super Dimension InfoTech Ltd, China), which mainly consists of 18 high-speed industrial cameras, each with 5 million pixels. The 3D model of the ultrasound patch on the rat’s skin was then reconstructed by the system, and the curvature of the ultrasound patch was calculated using an open source software (CloudCompare).

### Acoustic field measurements

The experimental setup of acoustic field measurement includes a water tank filled with degassed and deionized water, a three-axis stepper motor, an oscilloscope (DSOX3054G, Keysight, USA), a hydrophone (NH0200, Precision Acoustics, UK), a Vantage HIFU system and the ultrasound patch. The patch was attached to a sound-absorbing sponge and a 3D printed curved mold designed to match the curvature of the the patch on a rat’s skin (Supplementary Fig. 11). The Vantage system was used to apply electrical signals and time delays to the ultrasound patch. Data recorded by the oscilloscope were analyzed using MATLAB in accordance with the hydrophone calibration datasheet.

### Ultrasound thermal effect experiments

In the ultrasound thermal effect experiments, pork tissue embedded with thermocouples was placed at the focal length distance from the ultrasound patch. A sound-absorbing sponge was used as a flat surface, while two 3D-printed molds were utilized to create curved surfaces (Supplementary Fig. 11). A three-axis stepper motor was used to adjust the position of the patch. The focal point was determined by locating the position where the thermocouples recorded their maximum temperature peak. A software-controlled function generator switch was used to turn off the ultrasound stimulation after a duration of 5 minutes. All data were subsequently processed using MATLAB. During the experiments, both the tissue and the ultrasound patch were immersed in degassed and deionized water, with the water temperature maintained at a constant 37°C.

### Ultrasound treatment

In the single treatment and multiple treatment groups, the rats were first anesthetized with isoflurane. The location of the sciatic nerve was identified using a B-mode ultrasound probe, which then guided the placement of the ultrasound patch. The ultrasound patch was electrically connected to a waveform generator (Keysight 33500B, USA), set at a frequency of 1.5 MHz, an amplitude of 600 mVpp, 20 burst cycles, and an 80% duty cycle, along with a power amplifier (Model 100A400A, USA) set to 80% gain. Each rat underwent a 5-minute stimulation session a day.

### Behavioral tests

Behavioral tests were performed to assess the treatment effects, including a mechanical pain test and a thermal pain test. For the mechanical pain test in diabetic rats, the paw mechanical withdrawal threshold (PWT) was determined using the Von Frey method. Each rat was placed in a plastic box with a metal mesh floor for 20 minutes to acclimate them to the environment. Von Frey filaments (Aesthesio, USA) of varying forces (ranging from 0.16 to 26 g) were manually applied to stimulate the soles of the right hind paw. The assessment began with a 2 g filament, and the up-down method (SUDO) was employed to calculate the PWT. Behaviors such as lifting or licking the feet were considered positive reactions.

The thermal pain test for diabetic rats involved measuring paw thermal withdrawal latency (PWL) through the following procedure. Initially, each rat was placed on a hot plate set at 25°C for 10 minutes to acclimate to the environment. Subsequently, the rat was transferred to another hot plate maintained at 55°C. PWL was defined as the time elapsed from when the rat stood firmly until it exhibited one of the following behaviors: licking the hind paw, flicking the hind paw abnormally, or jumping to escape from the hot plate. All thermal pain test behavior experiments were recorded on video for further validation and to ensure a blind experimental design.

### Nerve conduction velocity test

Each rat was anesthetized with isoflurane before the compound action potential (CAP) of its sciatic nerve was recorded using an iWorx RA Recorder (IX-RA-834, iWorx Systems, USA). The anodic recording electrode was inserted at the rat’s ankle, while the cathodic recording electrode was placed proximally to the anodic recording electrode. A reference electrode was placed between the two recording electrodes. For stimulation, two electrodes were placed between the rat’s thighs and calves to deliver a 0.1 ms electrical pulse signal with a voltage amplitude of 5 V. The nerve conduction velocity was calculated by dividing the distance between the anodic stimulating electrode and the cathodic recording electrode by the CAP latency.

### Safety evaluation: Mechanical index, *I_spta_*, *I_sppa_*, and CEM43°C

The peak negative acoustic pressure *p* can be calculated from the voltage *v(t)* obtained from the hydrophone.

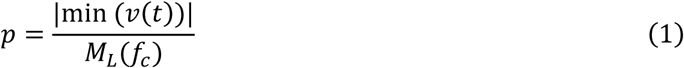

where *M*_*L*_(*f*_*c*_) is the hydrophone sensitivity conversion factor at the center frequency *f*_*c*_, and its unit is V/Pa.

And the mechanical index (MI) can be calculated from:

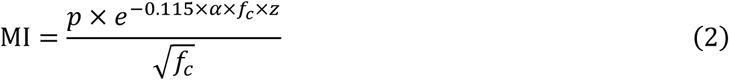

where *⍺* is the attenuation factor, which is 0.3 dB/(cm × MHz) in FDA, and z is the depth in cm.

The pulse integral intensity *PII* is calculated from:

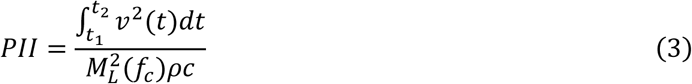

where *t*_1_ is the starting time of a single ultrasound pulse, measured in seconds, and *t*_2_ is the ending time of the same ultrasound pulse, measured in seconds. And *ρ* is the density of water (1000 kg/m^3^), and *c* is the speed of sound in water (1480 m/s). The spatial peak temporal average intensity (*I*_*spta*_) is calculated from:

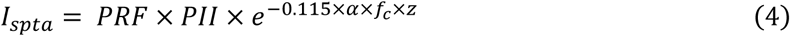

where PRF is the pulse repetition frequency, measured in Hz. And the pulse duration (*PD)* is calculated from:

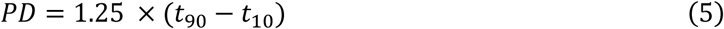

where *t*_10_ is the time when the amplitude is 10% below peak PII and *t*_90_ is the time when the amplitude is 90% below peak PII.

And the spatial peak pulse average intensity (*I*_*sppa*_) is calculated from:

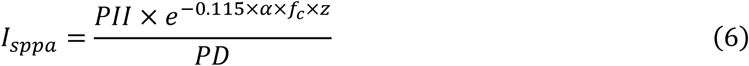

The cumulative equivalent minutes at 43 °C (CEM43°C) thermal dose was used to assess the safety of the ultrasound sequence [46], which is calculated by using the following formula:

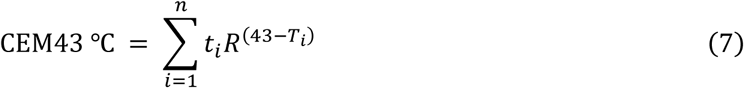

where CEM43°C represents the cumulative equivalent minutes at 43 °C, *t*_i_ denotes the *i*-th time interval, *R* is a factor related to the temperature-dependent rate of cell death (*R* = 0.25 if *T* < 43°C, *R* = 0.5 if *T* > 43 °C), and *T* represents the average temperature during time interval *t_i_.* The resulting CEM43°C value quantifies the overall impact of heat exposure on cell death. In clinical practice, a CEM43°C value below 2 min is generally considered safe for any type of tissue, provided that the procedure is monitored by a doctor or trained professional.

Temperature data for the patch and skin surface were obtained from the infrared images captured using a handheld thermal infrared imager (HM-TPH21Pro-3AQF, HIKMICRO). The temperature profile and thermal image of the w-SUP attached to the forearm are shown in Supplementary Fig. 8.

### Histological analysis

Histological analysis was performed to evaluate the safety of the focused ultrasound (FUS) used in the study. The rats were euthanized via cervical dislocation under anesthesia, and the FUS-treated sciatic nerves were harvested. These nerve samples were then fixed in 4% paraformaldehyde and subsequently stained with Hematoxylin and Eosin (H&E) by a technician who was blinded to the group assignments.

### Assessment of cell viability under ultrasound exposure

In the cell viability assay, five groups were studied: a control group treated with 75% ethanol for 10 min, and four experimental groups exposed to ultrasound for 0 min, 5 min, 15 min, and 30 min, respectively, with two samples per experiment. PC12 cells were cultivated in a specialized culture medium at 37°C, with 95% humidity, and 5% CO_2_. The cells were inoculated at a density of 1 ×10^4^ ml^−1^ into a 24-well cell culture plate and incubated for an additional 24 h. No antibodies were used in the experiment. Ultrasound exposure was applied to the bottom of the culture plate for 5, 15, and 30 min. Following the exposure, cells were stained with calcium green AM (excitation/emission = 488 nm/525 nm) and propidium iodide (excitation/emission = 530 nm/620 nm) for 15 min. The cells were then imaged using a fluorescence microscope. The positive control group consisted of cells treated with 75% ethanol for 10 min.

### Statistical analysis

All experimental data are presented as mean ± standard deviation (SD) and analyzed using Prism software (GraphPad Prism, GraphPad Software, USA). Two-way ANOVA followed by Tukey’s multiple comparisons was performed to compare paw withdrawal responses across different groups. In these tests, p-value of <0.05*, <0.01**, <0.001***, <0.0001**** were considered statistically significance.

## Data availability

The data that support the findings of this study have been included in the main text and Supplementary Information. All other relevant data supporting the findings of this study are available from the corresponding authors upon request.

## Acknowledgements

This work was supported by the National Science Foundation of China (82151318 and 12204307) and the Shanghai Sailing Program (22YF1428900).

## Author contributions

C. Peng conceived the research idea and directed all research activities. B. Fu designed the device and performed simulaitons. C. Pu and B. Fu fabricated and characterized the device, conducted experiments and wrote the manuscript. X. Guan helped with the rats experiments and biological safety test. P. Lu executed the rats experiments. J. Zhang measured the curvature of the device on a rat’s skin. Y. Shen, X. Li, and L. Guo helped with the rats experiments. H. Xu and X. Jiang assisted with manuscript preparation. All authors contributed to manuscript writing.

## Competing interests

The authors declare no competing interests.

## Additional information

None

## Notes

### Competing Interest Statement

The authors have declared no competing interest.

