## Supplementary information for "A Wearable Spiral Ultrasound Patch for Focused Ultrasound Peripheral Neuromodulation of Diabetic Neuropathic Pain"

### Contents

#### Supplementary Figures

Supplementary Fig. 1 | Optical image and results of cyclic tensile tests.

Supplementary Fig. 2 | Simulated acoustic beam profiles of transducers in square, circular and spiral patterns.

Supplementary Fig. 3 | Simulated acoustic beam profiles of transducers in a spiral pattern at different frequencies.

Supplementary Fig. 4 | Frequency-impedance and frequency-phase spectra of the four channels of the w-SUP before impedance matching.

Supplementary Fig. 5 | Measured acoustic pressure at the focal point of the w-SUP adhered to three different surface types.

Supplementary Fig. 6 | Optical image and the captured image of the w-SUP conforming to a rat's skin.

Supplementary Fig. 7 | Schematic diagram illustrating the treatment processes for the single treatment and multiple treatments groups.

Supplementary Fig. 8 | Thermal impact of the w-SUP on the forearm.

Supplementary Fig. 9 | Optical images of the top and bottom electrodes.

Supplementary Fig. 10 | Schematic diagrams illustrating the fabrication process.

Supplementary Fig. 11 | Schematic diagrams of the curved surfaces with 20° and 40° angles.

#### Supplementary Tables

Supplementary Tab. 1 | Simulated acoustic pressure at the focal point of transducer arranged in circular and spiral patterns.

Supplementary Tab. 2 | Measured full width at half maxima (FWHM) in the Y-X and Z-X planes of the w-SUP.

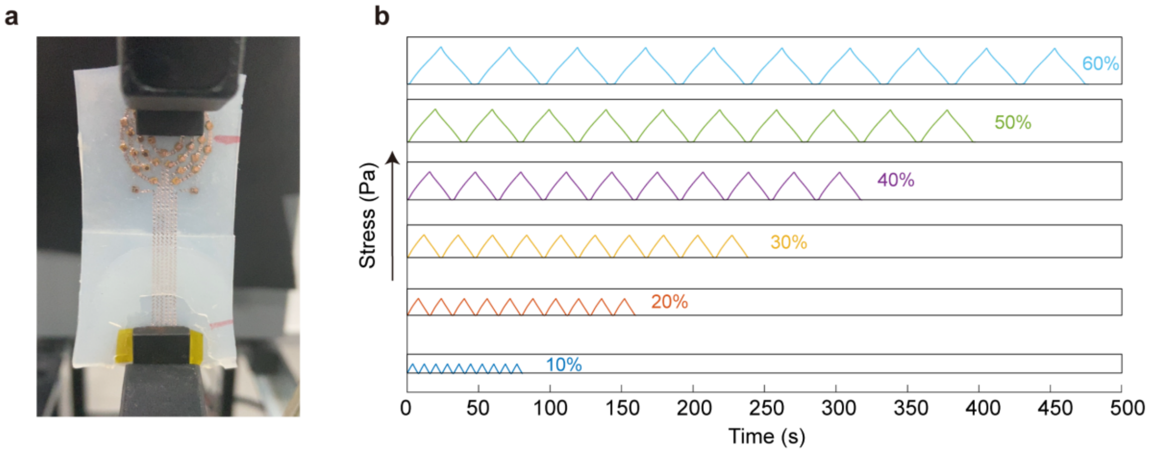


**Supplementary Fig. 1 | Optical image and results of cyclic tensile tests. a** Optical image of the tensile performance test. **b** Cyclic tensile testing curve, displaying strain ranges from 10% to 60% over 10 cycles per test.

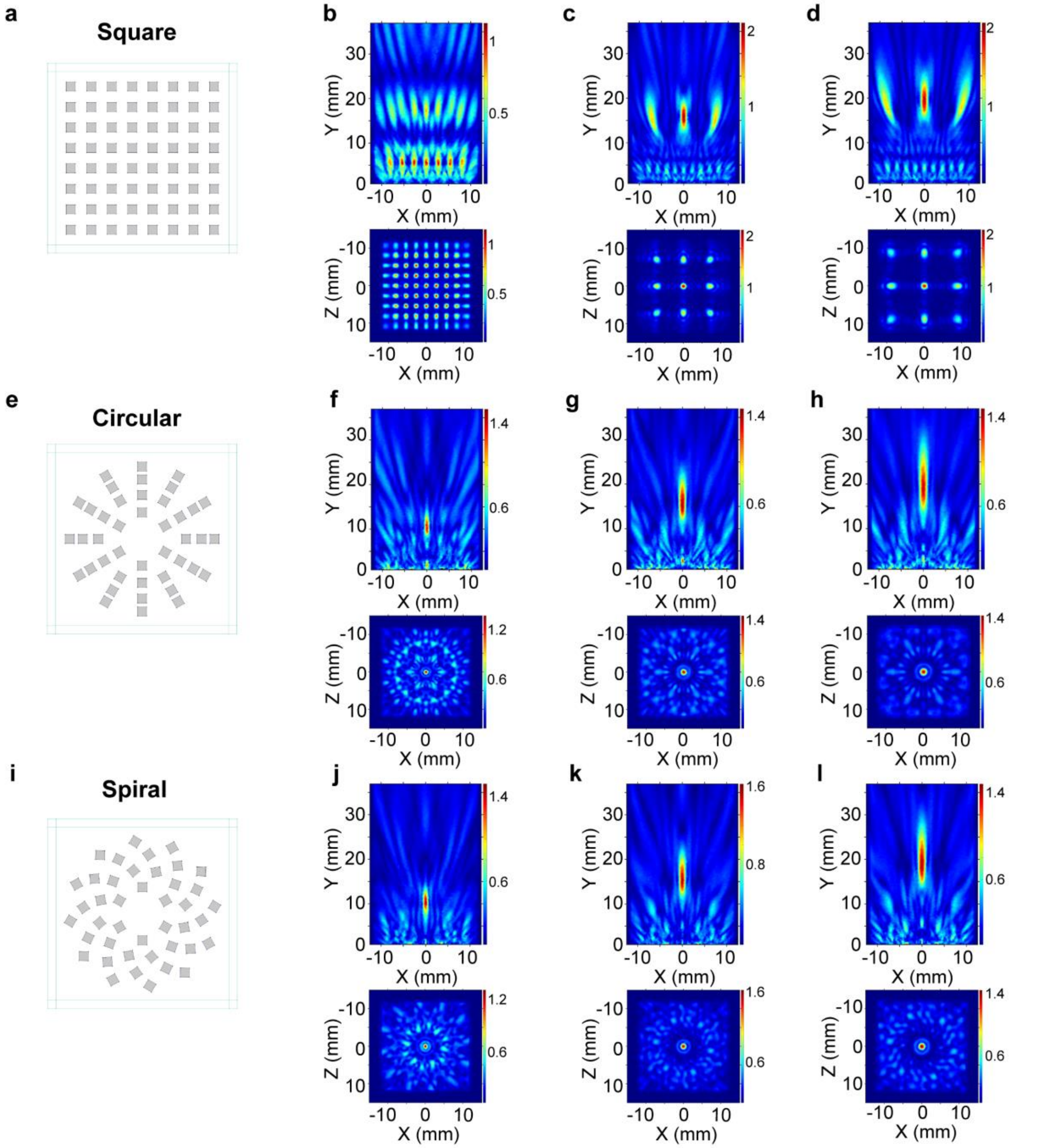


**Supplementary Fig. 2 | Simulated acoustic beam profiles of transducers in square, circular and spiral patterns.** **a** Schematic of transducer elements arranged in a square matrix pattern. Simulated acoustic beam profiles in the Y-X axis and Z-X axis of the square pattern without phase control (**b**), and with targeted focal lengths of 14 mm (**c**), and 20 mm (**d**). **e** Schematic of transducer elements arranged in a circular pattern. Simulated acoustic beam profiles in the Y-X axis and Z-X axis of the circular pattern without phase control (**f**) with targeted focal lengths of 10 mm, 14 mm (**g**), and 20 mm (**h**). **i** Schematic of transducer elements arranged in a spiral pattern. Simulated acoustic beam profiles in the Y-X axis and Z-X axis of the spiral pattern without phase control (**j**), with targeted focal lengths of 14 mm (**k**), and 20 mm (**l**). The simulated frequencies in **b**-**d, f**-**h,** and **j**-**l** are all set at 1.5 MHz.

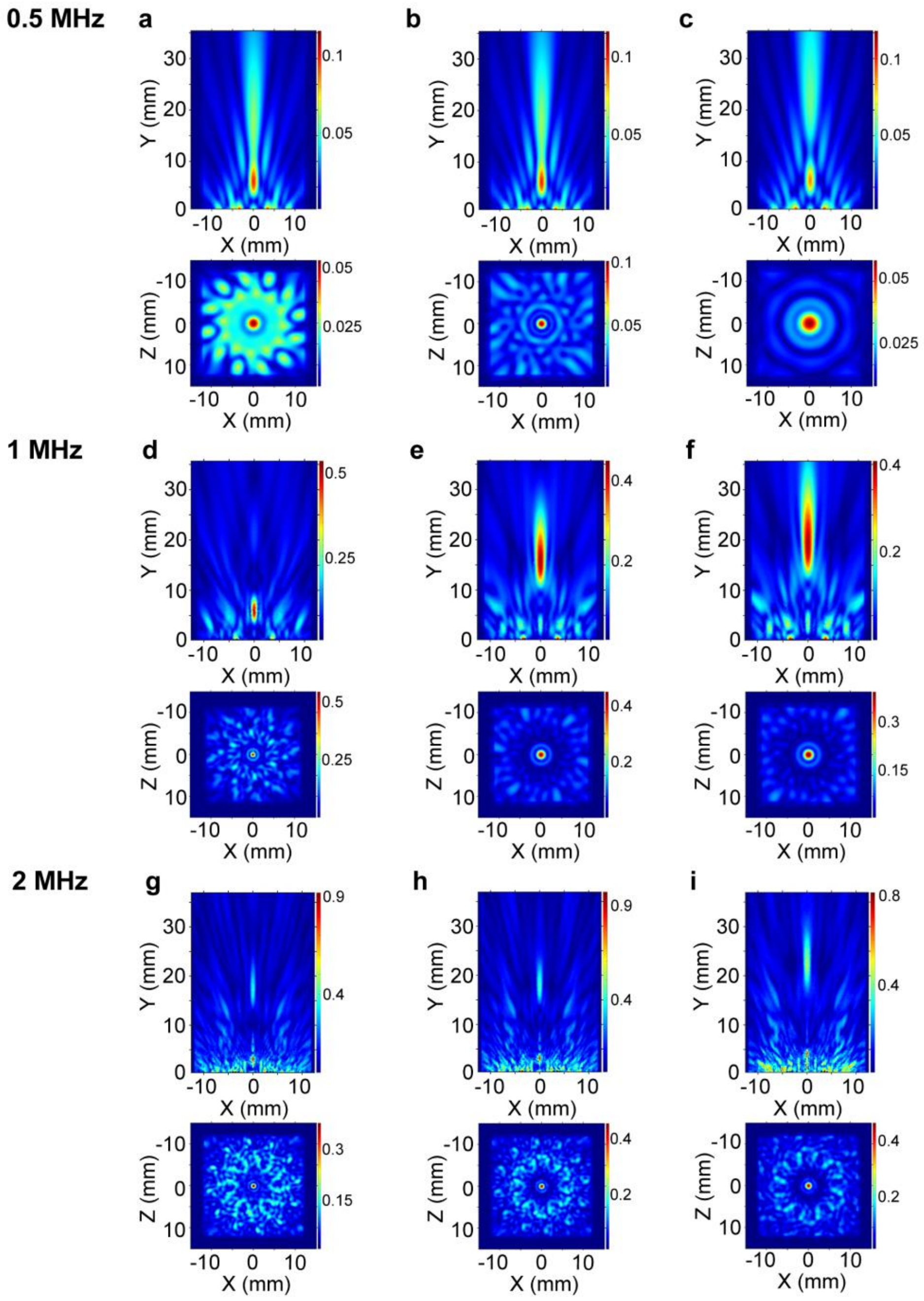


**Supplementary Fig. 3 | Simulated acoustic beam profiles of transducers in a spiral pattern at different frequencies.** Simulated acoustic beam profiles in the Y-X axis and Z-X axis of the spiral pattern at a frequency of 0.5 MHz, without phase control (**a**), and with targeted focal lengths of 14 mm (**b**) and 20 mm (**c**). Simulated acoustic beam profiles in the Y-X axis and Z-X axis of the spiral pattern at a frequency of 1 MHz, without phase control (**d**), and with targeted focal lengths of 14 mm (**e**), and 20 mm (**f**). Simulated acoustic beam profiles in the Y-X axis and Z-X axis of the spiral pattern at a frequency of 2 MHz, without phase control (**g**), and with targeted focal lengths of 14 mm (**h**), and 20 mm (**i**).

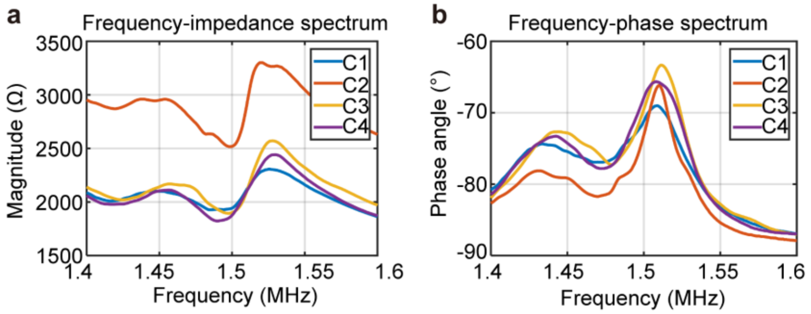


**Supplementary Fig. 4 | Frequency-impedance and frequency-phase spectra of the four channels of the w-SUP before impedance matching. a** Frequency-impedance spectrum of the four channels of the w-SUP prior to impedance matching. **b** Frequency-phase spectrum of the four channels of the w-SUP prior to impedance matching.

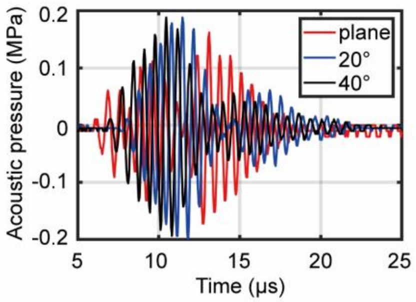


**Supplementary Fig. 5 | Measured acoustic pressure at the focal point of the w-SUP adhered to three different surface types: red indicates a flat surface, blue represents a 20° curved surface, and black signifies a 40° curved surface.**

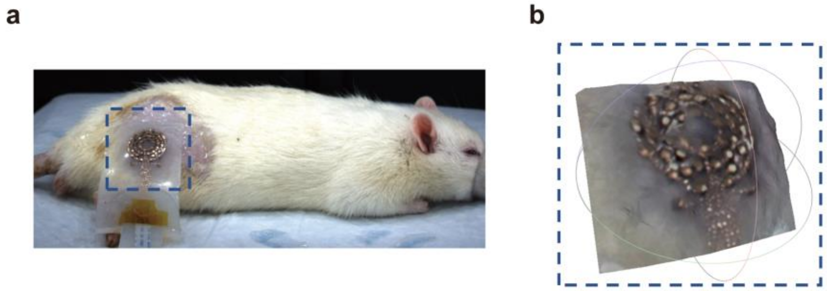


**Supplementary Fig. 6 | Optical image and the captured image of the w-SUP conforming to a rat's skin. a** Optical image of the w-SUP attached to a rat's skin. **b** Captured image of the w-SUP attached to a rat's skin with a radius curvature of 18.34 cm.

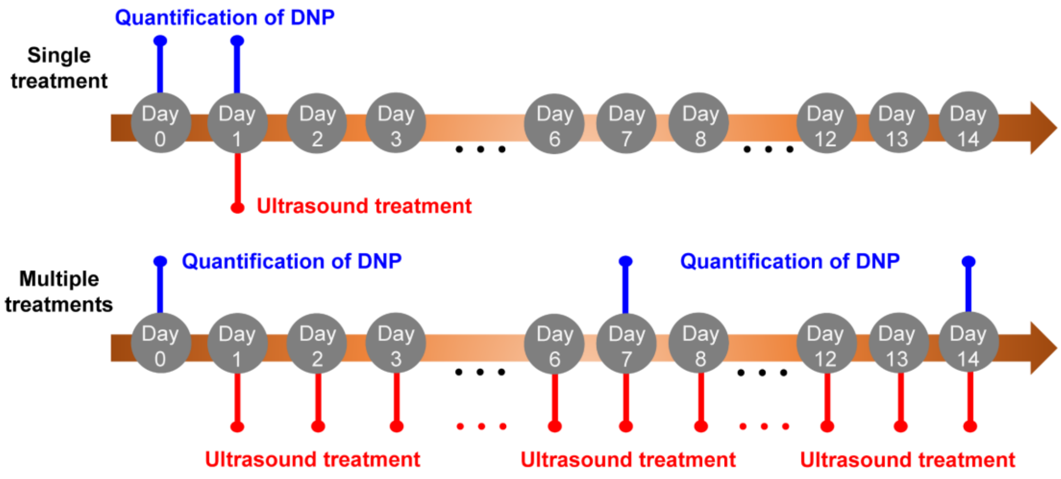


**Supplementary Fig. 7 | Schematic diagram illustrating the treatment processes for the single treatment and multiple treatments groups.**

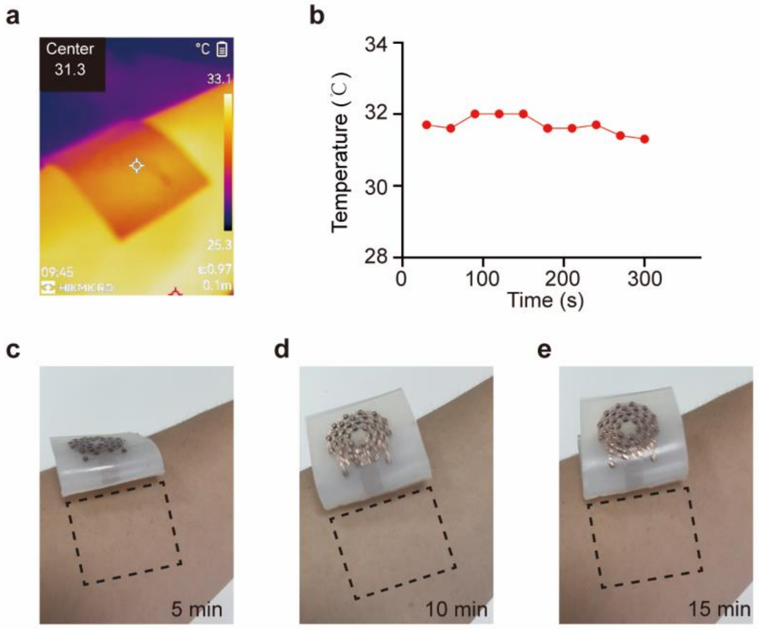


**Supplementary Fig. 8 | Thermal impact of the w-SUP on the forearm. a** Thermal IR image of the ultrasound patch applied to the forearm. **b** Skin temperature profile recorded during a 5-minute ultrasound application to the forearm. **c-e** Optical images depicting the skin condition after the w-SUP was attached for 5, 10, and 15 minutes, respectively.

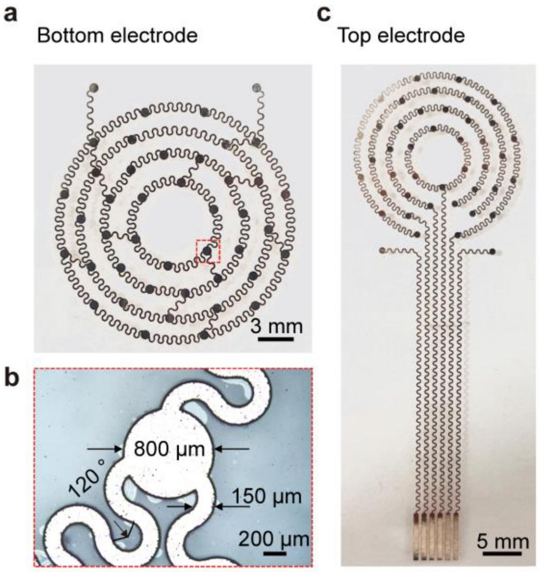


**Supplementary Fig. 9 | Optical images of the top and bottom electrodes. a** Optical image of the bottom electrode. **b** Close-up view of the section highlighted by the red box in (a). **c** Optical image of the top electrode.

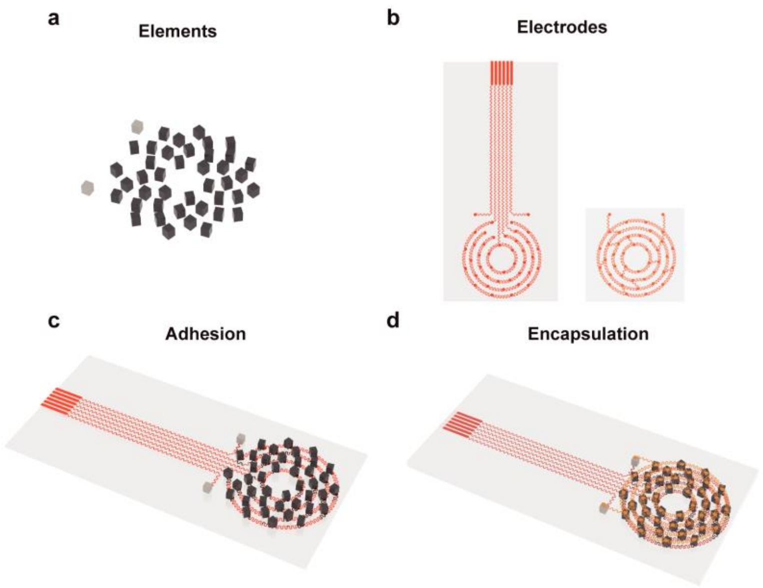


**Supplementary Fig. 10 | Schematic diagrams illustrating the fabrication process. a** Schematic diagram of 41 PZT-4 transducers and 2 VIAs (Vertical interconnect access). **b** Schematic layout showing the top and bottom electrodes embedded with the Ecoflex substrate. **c** Schematic diagram of the adhesion between the transducer elements and the top electrode. **d** Schematic diagram of the ultrasound patch after Ecoflex encapsulation.

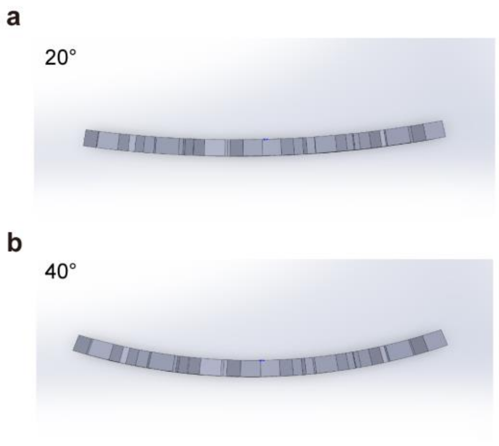


**Supplementary Fig. 11 | Schematic diagrams of the curved surfaces with 20° and 40° angles. a** Schematic diagram of the 20° curved surface with a radius of curvature of 18.34 cm. **b** Schematic diagram of the 40° curved surface with a radius of curvature of 9.17 cm.

**Supplementary Tab. 1 | Simulated acoustic pressure at the focal point of transducer arranged in circular and spiral patterns.**

| Targeted focal length (mm) | Simulated acoustic pressure at focal point of transducer in circular pattern (MPa) | Simulated acoustic pressure at focal point of transducer in spiral pattern (MPa) |
|---|---|---|
| Without phase control | 1.36 | 1.48 |
| 14 | 1.43 | 1.60 |
| 20 | 1.42 | 1.43 |

**Supplementary Tab. 2 | Measured full width at half maxima (FWHM) in the Y-X and Z-X planes of the w-SUP.**

| Measured focal length (mm) | Targeted focal length (mm) | FWHM in the Y-X plane (mm) | FWHM in the Z-X plane (mm) |
|---|---|---|---|
| 11 | Without phase control | 10.4 | 1 |
| 12.7 | 14 | 11.8 | 1.2 |
| 17.5 | 20 | 14.5 | 1.6 |
